# Molecular dynamics analyses of CLDN15 pore size and charge selectivity

**DOI:** 10.1101/2023.08.16.553400

**Authors:** Sarah McGuinness, Pan Li, Ye Li, Shadi Fuladi, Sukanya Konar, Samaneh Sajjadi, Mohammed Sidahmed, Yueying Li, Le Shen, Fatemeh Khalili Araghi, Christopher R. Weber

## Abstract

Claudin-15 (CLDN15) molecules form channels that directly regulate cation and water transport. In the gastrointestinal tract, this transport indirectly impacts nutrient absorption. However, the mechanisms governing ion transport through these channels remain poorly understood. We addressed this question by building on our previous cell culture studies and all atom molecular dynamic simulation model of CLDN15. By mutating D55 to a bulkier glutamic acid (E) or neutral amino acid asparagine (N), our *in vitro* measurements showed that the D55E mutation decreased charge selectivity and favored small ion permeability, while the D55N mutation led to reduced charge selectivity without markedly altering size selectivity. By establishing a simplified (reduced) CLDN15 molecular dynamics model that excludes non-essential transmembrane regions, we were able to probe how D55 modified cation dehydration, charge interaction, and permeability. These results provide novel insight into organization of the CLDN15 selectivity filter and suggests that D55 plays a dual role in shaping both electrostatic and steric properties of the pore, but its electrostatic role is more prominent in determining CLDN15 cation permeability. This knowledge can be used toward the development of effective strategies to modulate CLDN15 function. The experimental approach established can be further extended to study the function of other claudin channels. Together, these advancements will help us to modulate tight junctions to promote human health.

**SUMMARY:** Cell culture and molecular dynamics simulations reveal the role of the CLDN15-D55 residue in ion size and charge selectivity. A reduced CLDN15 model offers novel insights into ion conductance, providing a valuable tool for therapeutic modulation of tight junctions.

## INTRODUCTION

The claudin (CLDN) family of tight junction proteins plays critical roles in defining paracellular permeability. Tight junction strands containing claudin proteins form circumferential meshworks at the apical end of the intercellular space by interacting in cis- and trans- at the cellular membrane(Krause et al., 2015). Claudins within this meshwork are critical in regulating paracellular passage of ions, solute, and water across epithelial tissues(Furuse et al., 1998; Morita et al., 1999; Furuse, 2010). Members of the CLDN family have diverse effects on tight junction barrier function, with some forming ion-permeable pathways and others restricting ion flux(Van Itallie et al., 2003; Piontek et al., 2011). Channel-forming CLDNs, such as CLDN15, CLDN10a/b, and CLDN2 selectively regulate paracellular flux of molecules in a size and charge selective manner (Amasheh et al., 2002; Colegio et al., 2002; Günzel et al., 2009; Yu, 2009; Krug et al., 2012; Rosenthal, 2017; Rosenthal et al., 2020; Gonschior et al., 2022; Hempel et al., 2022). For example, in the mammalian intestine, CLDN15 is critical to the paracellular permeability of monovalent cations such as Na^+^, which provides a driving force for Na^+^-dependent nutrient absorption (Tamura et al., 2011).

Structural differences between claudins can significantly influence ionic interactions within the channel, affecting small molecule permeability in a size- and charge-selective manner(Krause et al., 2009; Piontek et al., 2011; Krause et al., 2015; Piontek, 2017). CLDN15 was the first member of the family to have a partially resolved 3D-structure, leading to a proposed model for claudin strands featuring two antiparallel double rows of claudins in opposing lipid bilayers(Hiroshi Suzuki 2014; Hiroshi Suzuki, 2015). Subsequent molecular dynamics simulations by us and others refined this atomic model of CLDN15 paracellular ion channels (Alberini et al., 2017, 2018; Samanta et al., 2018). These computational studies revealed the detailed structure of the pore and the site of CLDN15’s selectivity filter: two rings of aspartic acid residues (D55) near the center of the pore generate a negative electrostatic potential to encourage cation flux (Colegio et al., 2002; Colegio, 2003; Van Itallie et al., 2003; Angelow et al., 2008; Samanta et al., 2018; Hempel et al., 2022).

Understanding how claudin structure regulates paracellular flux is central for comprehending extracellular fluid homeostasis in different epithelia, and for determining the effects of altered claudin expression or mutations. To further investigate the ion transport mechanisms of CLDN15, we developed a reduced model of CLDN15 channel that allowed us to extend simulation durations while preserving key features of ion diffusion. This model enabled efficient simulations of multiple mutants under varying conditions such as cation size and salt concentration.

We found that mutating D55 to glutamic acid (E) or asparagine (N) altered pore selectivity, providing insights into the mechanisms of ion diffusion through CLDN15. These mutations significantly affected charge selectivity in both *in vitro* and *in silico* studies, while also revealing subtle size-selectivity differences that were most clearly detected *in vitro*. These results illustrate the nuanced interplay between size and charge selectivity within CLDN15, regulated by a single key residue in the selectivity filter. The ability of D55 to control both electrostatic interactions and cation dehydration highlights a dual mechanism for defining the channel’s permeability. The reduced CLDN15 model we present will be a valuable tool for future studies aimed at regulating claudin channel function, with potential applications in modulating tight junction permeability in health and disease.

## MATERIALS AND METHODS

### Generation of MDCK cells with inducible expression of CLDN15 and CLDN15 pore mutants

**C**LDN15^WT^, CLDN15^D55E^ and CLDN15^D55N^ expressing constructs were made similarly to previously described (Quintara Biosciences, Hayward, CA), (Samanta et al., 2018) , and stably transfected into MDCK I cells along with a *PiggyBac* transposase encoding plasmid. After hygromycin selection, pooled clones were used for subsequent experiments.

### Barrier function measurements in tissue cultured epithelial monolayers

MDCK I Cells were plated on 0.33 cm^2^ polycarbonate semipermeable Transwell supports with 0.4 µm pores (Corning Life Sciences) at confluent density as previously described (Samanta et al., 2018). CLDN15 expression was induced using doxycycline (10–50 ng/ml) 1 d after plating. Doxycycline dosage was chosen to generate matched CLDN15 expression, as assessed by fluorescence microscopy. At day 5, cells were transferred to HBSS with 1 g/ml glucose and incubated at 37°C for 30 min prior to measurement of transepithelial electrical resistance (TER). Measured TER values were corrected for the resistance of the system in the absence of cells. Reversal potential measurements were performed using current clamp to I=0 in the setting of asymmetrical apical: basolateral ionic compositions. For NaCl dilution potentials, reversal potentials were measured with basal HBSS replaced by HBSS containing half the apical NaCl concentration (osmotically balanced with mannitol). From the reversal potential, the Goldmann–Hodgkin–Katz (GHK) equation was used to calculate the relative permeability of sodium to chloride (V_rev_ = −(RT/F)ln[(α + β)/(1 + αβ)], where β = PCl^−^/PNa^+^, α = 1.9, RT/F = 26.6). For biionic potentials, reversal potentials were measured with Na^+^ of basal HBSS replaced by different monovalent cations. The GHK equation was used to calculate the permeability of different sized cations relative to Na^+^ (V_rev_ = (RT/F) ln[(γ + β)/(1 + β)], where γ = PM^+^/PNa^+^) (Yu et al., 2009).

### Molecular dynamics simulations

Molecular dynamics (MD) simulations were performed using the program NAMD (Phillips et al., 2005; Phillips et al., 2020) We used the CHARMM36 force field for proteins (MacKerell, 1998; Mackerell Jr, 2004; Best, 2012), ions (Beglov and Roux, 1994), phospholipids (Klauda et al., 2010), and TIP3 water model (Jorgensen, 1983). The updated nonbonded interactions between monovalent cations for their interaction with Cl^-^ were taken from (Yoo and Aksimentiev, 2012) and applied through NBFIX correction in the forcefield. The forcefield parameters for methylammonium (MA^+^), ethylammonium (EA^+^), tetramethylammonium (TMA^+^), and tetraethylammonium (TEA^+^) were taken from the CHARMM General ForceField (CGenFF). All simulations were carried out with periodic boundary conditions and a time step of 1 fs. Simulations were performed at constant temperature (333K) using Langevin dynamics with a friction coefficient of ϒ = 5ps^-1^, and pressure was maintained at 1 atm using the Langevin Nosé-Hoover method (Feller et al., 1995). We calculated the long-range electrostatic forces using the particle mesh Ewald (PME) algorithm with a grid spacing of at least 1 Å in each direction (Darden et al., 1993). A cutoff of 12 Å was used for non-bonded (short-range electrostatic and VdW) interactions.

### System setup of CLDN15 (full channel)

A full model of CLDN15 channels in two parallel POPC lipid bilayers was constructed based on our previously refined model of CLDN15 pores (Samanta et al., 2018). The unit cell in this model consists of 12 CLDN15 monomers forming three ion channels defining conductance in a plane parallel to the lipid bilayer. The entire system was comprised of 341,000 atoms.

To ensure the homogeneity of the system, the equilibrated conformation of the middle tetrameric pore was replaced by one of the other to the other pores, and the system was further equilibrated as follows: The lipid and water molecules were unaltered, and the system was ionized with NaCl at 180 mM. The charges across the membrane were balanced by adding 24 additional Na^+^ ions to the middle compartment to balance out the negative charge of the protein. During equilibration, the protein backbone was initially constrained with a force constant of 1.0 kcal/mol·Å^2^ and then gradually released to 0.75, to 0.50, to 0.25 and to 0.10 kcal/mol·Å^2^ over a period of 30 ns. The constraints were then released for 10 ns, during which all three CLDN15 channels remained stable with an RMSD comparable to trajectories of our CLDN15 model published previously(Samanta et al., 2018).

Eight additional systems corresponding to LiCl, KCl, RbCl, CsCl, MACl, EACl, TMACl, and TEACl were prepared by replacing Na^+^ ions in the equilibrated full model (described above). Each new system was equilibrated for 5 ns at constant pressure to allow re-orientation of solvent around the ions.

### Reduced model of CLDN15

We developed a reduced model of CLDN15 pores based on our previously published 3-pore model of CLDN15(Samanta et al., 2018). In this model, part of the transmembrane (TM) helices of the proteins anchoring the pores to the membranes and their surrounding lipid membranes were omitted, leaving the ion conduction pore and its environment intact. The reduced model consists of the ion conduction pores and their surrounding solution with only 59,000 atoms.

To generate the reduced model, sections of the four TM helices enclosed by the lipid bilayer (CLDN15 residues 21 to 83 and 131 to 167) and the surrounding lipid membranes were removed. Water molecules between the two lipid bilayers were kept avoiding any changes in the ion conduction pores and their immediate surroundings. To emulate the anchoring role of the lipid bilayers, the four truncated TM helices were restrained in place by applying a harmonic constraint with a force constant of 5.0, 2.0, or 0.5 kcal/mol·Å^2^ to the backbone of the last three helical turns of the reduced model (residues 21-26, 77-83, 131-136, and 161-167). Water molecules were constrained to the initial slab by applying a flat-bottom harmonic potential of 10.0 kcal/mol·Å^2^ normal to the plane of membrane. The slab boundaries were chosen to match the equilibrated configuration, with a thickness of 30 Å on the two sides of the pore and a thickness of 42 Å in the middle of two rows. To avoid strong electrostatic interactions of the proteins with their periodic images, a periodic boundary was set in z-direction (normal to the membranes) 15 Å beyond the system.

This reduced model system was equilibrated for 105 ns with no additional restraints keeping the unit cell volume constant to assess its stability. This baseline system had an ion concentration of 300 mM.

Additional systems with reduced NaCl concentrations of 150 mM and 100 mM were prepared by removing ions to reach the desired concentration and equilibrated for 20 ns. During the first 10ns the protein backbone was constrained. Additional systems containing LiCl, KCl, RbCl, CsCl, MACl, EACl, TMACl, and TEACl in place of NaCl were prepared and equilibrated for 5 ns to ensure proper hydration of ions and water reorientation prior to application of a voltage bias. Cations located inside the pore were moved to the bulk solution before replacing Na^+^ with heavy cations to remove any bias for the low-conducting cations.

### Mutant channels

Two CLDN15 mutants, CLDN15^D55E^ and CLDN15^D55N^, were constructed from the reduced model system. Starting from the CLDN15^WT^ reduced model after 105 ns of equilibration, the two mutants were generated using VMD (Humphrey et al., 1996) to convert residue D55 on to glutamic acid (E) or asparagine (N). The systems were ionized to 100mM and neutralized by adjusting the number of Cl^-^ ions and were equilibrated before applying a voltage bias. The protein backbone was constrained for the first 13 ns of equilibration, followed by 20 ns of unrestrained equilibration.

These NaCl mutant systems were then used to construct eight other systems by replacing NaCl with LiCl, KCl, RbCl, CsCl, MACl, EACl, TMACl, or TEACl. To equilibrate these systems, we performed 2000 steps of minimization, 10 ns of 1.0 kcal/mol·Å^2^ backbone constraint, and finally 10 ns equilibration without backbone constraints.

### Ion transport simulations

The full model (WT) and the reduced model (WT and two mutants) systems were simulated under a voltage bias by application of a constant electric field along the channel’s axis to study ion transport (Aksimentiev et al., 2004; Aksimentiev and Schulten, 2005; Khalili-Araghi et al., 2013; Samanta et al., 2018). Due to the continuous setup of claudin pores across the periodic boundaries, a constant electric field will result in uniform voltage bias across claudin pores without any additional leaks (voltage drops). This enables us to simulate the ionic currents under a constant voltage and thus obtain the current-voltage (I-V) relationship of the channel and calculate the channel conductance.

The WT full model system was simulated with NaCl, LiCl, KCl, RbCl, CsCl, MACl, EACl, TMACl, and TEACl for 60 ns at ±100 mV, ±200 mV, and ±400 mV. The reduced model system at 150 mM salt concentration was simulated at ±200 mV, and ±400 mV for 70 ns each and was used as the basis of comparison with the full model. To prevent protein drift during simulations of ion transport, the backbone of the transmembrane (TM) helices anchoring the pores to the membrane in the full model were constrained with a force constant of 0.5 kcal/mol·Å^2^ when applying a voltage bias.

Two copies (replicas) of the reduced model (WT and two mutants) at 100 mM salt concentration were prepared and simulated under an applied voltage bias to calculate the ion conductance of WT and mutants. Each system was prepared from coordinates obtained from initial 105 ns equilibration of the reduced model, and each replica was initialized from random distribution of the ions. After ion placement, each system was equilibrated for 10 ns with protein backbone constraints (with a force constant of 1.0 kcal/mol·Å^2^), followed by 10 ns of equilibration without any constraints. These low salt concentration systems match the experimental conditions, and their trajectories were used for all the analysis reported here.

CLDN15^WT^, CLDN15^D55N^, and CLDN15^D55E^ were simulated with NaCl, LiCl, KCl, RbCl, CsCl, MACl, EACl, TMACl, and TEACl at ±800 mV and ±1200 mV for 140 ns each (3 systems x 9 salt solutions x 4 voltages x 140ns each x 2 replicas). The ionic current was calculated from the displacement charge of the ions over time at each voltage(Gumbart et al., 2012; Khalili-Araghi et al., 2013). The conductance was calculated as G=I/V.

### Contact time and channel occupancy

The contact time of the ions with each amino acid residue within the channel was calculated from 40 ns of voltage simulations (±800 mV and ±1200 mV). Residues within 4 Å of the ions were selected, and the occupancy of each residue was calculated as the average number of ions in proximity of that residue. These trajectories were also analyzed to estimate the average occupancy of the ions per channel for each alkali cation system. Anions and cations residing with the channel were counted and averaged over simulation time.

### Pore radius

The middle pore radius of each reduced model was calculated using the program HOLE (Smart et al., 1996) with MDAnalysis, resulting in time-averaged radius profiles. The center point was measured from the averaged coordinates of the selectivity filter, consisting of four D55 residues at the center of the pore. 650 frames from the CLDN15^WT^, CLDN15^D55E^, CLDN15^D55N^ voltage runs (140ns at - 800 mV) were used for these calculations. The pore radius was calculated and averaged over all three of the channels in the system. We chose to calculate the pore radius for the voltage simulations, because these were the longest trajectories available for all three channels.

The calculations were repeated for positive and negative voltages (±800 mV and ±1200 mV) of CLDN15^WT^ and for all the three pores in the unit cell. The results (not shown here) indicated that voltage has no effect on the pore radius, and thus, we chose to present the radius profile for only one of these runs. In addition, visual inspection of the pores and the RMSD and RMSF of protein suggest that application of the voltage does not affect the pore radius.

### Cation hydration shell

Hydration shell was quantified by counting water molecules within the first hydration shell of permeating ions. A cutoff of 3.5 Å was used for Na^+^ and Li^+^ with ionic radii of 0.95 Å and 0.60 Å, respectively. For larger cations, a cutoff of 4.0 Å is sufficient to capture the hydration shell.

For CLDN15^WT^, CLDN15^D55E^, and CLDN15^D55N^ reduced model systems, the last 40 ns of 140ns (±800 mV and ±1200 mV) voltage trajectories were analyzed to calculate the average number of coordinating water molecules in the hydration shells of alkali cations.

### Cation forcefield parameters (bulk conductivity)

To validate the ion permeabilities calculated from simulations and the effect of forcefields and its shortcomings, we calculated the bulk conductivity of ions from molecular dynamics simulations and compared them against experiments (Table 1).

**Table 1.**
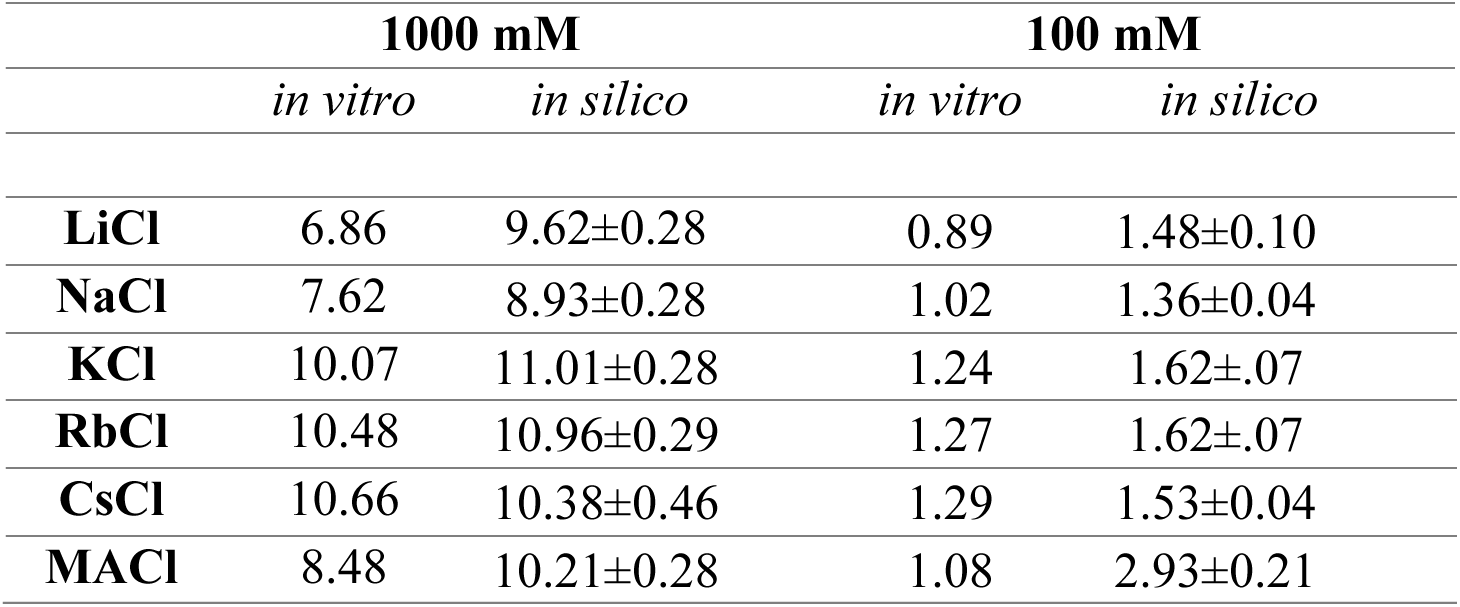
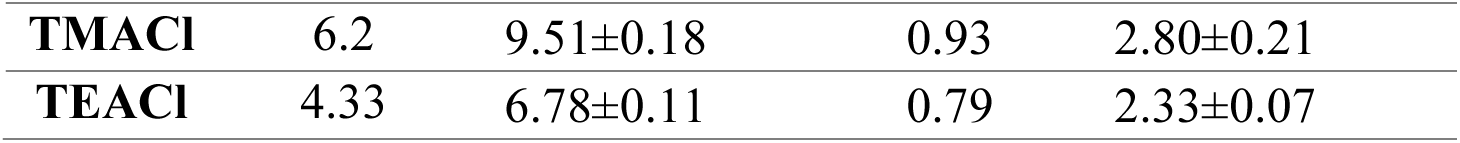
*In vitro* and *in silico* electrolyte solution bulk conductivity in Siemens/meter. *In silico* values are represented by average and standard deviation (SD).

For the 100 mM simulations, we generated a 100 × 100 × 100 Å^3^ pre-equilibrated water box containing 63 cations, 63 Cl^-^ ions and 96,000 total atoms. For 1000 mM simulations, we generated a 50 × 50 × 50 Å^3^ pre-equilibrated water box containing 75 cations and 75 Cl^-^ ions and 11,000 atoms. The system was equilibrated for 1 ns before applying a voltage bias of ±200 mV for 15 ns (constant volume, T=333 K) to generate a current. The last 11 ns of the simulation trajectory was used to determine the displacement charge and the solution conductivity similar to the other simulations reported here. The errors were calculated as standard deviation of the conductivities among multiple voltages. We also measured bulk conductivity of the same ions by determining the I-V relationships of 100 mM X^+^Cl^-^solutions filled in polyethelene tubes. Conductance was normalized by tube length and cross-sectional area.

The simulated and measured conductivity values were similar, as determined by linear regression for solutions of LiCl, NaCl, KCl, RbCl, CsCl, MACl, TMACl (slope=1.21, R^2^=0.99 at 100 mM and slope=1.06, R^2^=0.94 at 1 M). This indicates that the forcefield parameters used in the simulations are reliable and provide a good approximation of the real-world behavior of the cations in this study regardless of the salt concentration.

## RESULTS

We previously developed an all-atom molecular dynamic model for CLDN15 quaternary structure forming cation-selective ion channels (Samanta et al., 2018; Fuladi et al., 2022a; Fuladi et al., 2022b) In this model, claudin monomers possess four transmembrane (TM) helices and two extracellular segments (ECS1 and ECS2). Claudin monomers assemble into an antiparallel double row of claudins in lipid membrane, with extracellular β-sheets forming a half-pipe structure maintained by hydrogen bonds between β-strands (Suzuki, 2015, 2014). The pores are formed by two opposing half-pipes from two membranes and are sealed by hydrophobic trans-claudin interactions. The model, along with our cell culture studies, suggests the importance of pore-lining residue D55 as a principal determinant of channel selectivity (Alberini et al., 2017, 2018; Samanta et al., 2018).

### CLDN15 D55 is a critical determinant of pore function

To define the potential electrostatic versus steric determinants of the key residue D55 on pore permeability properties, we developed MDCK cell lines with tet-inducible expression of wild-type CLDN15 (D55) or two different pore mutants. For the first mutant, aspartic acid was changed to similarly charged but larger glutamic acid (D55E). For the second mutant, aspartic acid was changed to similarly sized, but electroneutral asparagine (D55N). These mutations were chosen to evaluate the potential contributions of size and charge of amino acid residue responsible for the permeability filter (D55) to claudin-15 pore permeability.

Immunofluorescent staining revealed appropriate and similar expressions of CLDN15^WT^, CLDN15^D55E^, and CLDN15^D55N^ at the tight junction (Fig. 1A). Expression of CLDN15 and its mutants significantly reduced transepithelial electrical resistance (TER; Fig. 1B). We next determined relative permeabilities of Na^+^ to Cl^-^ in these CLDN15 expressing monolayers by performing dilution potential measurements. The charge-maintaining D55E mutant expressing monolayers reduced PNa+/PCl- to 45% of that in CLDN15^WT^ expressing monolayers. More dramatically, the charge-neutralizing D55N mutant reduced PNa+/PCl- to 24% of CLDN15WT expressing monolayers (Fig. 1C).

**Figure 1:**
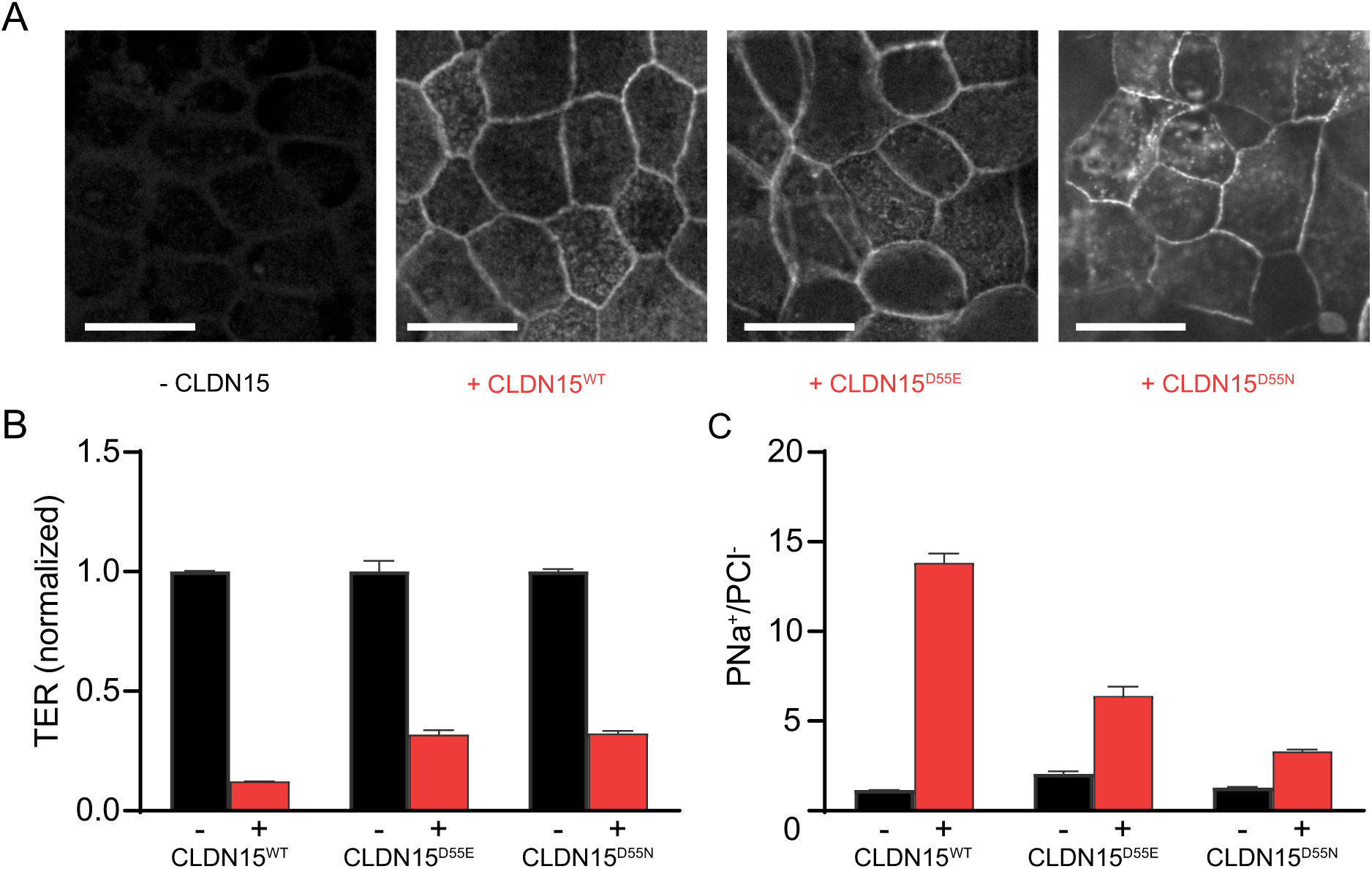
Generation and initial characterization of tet-inducible CLDN15-expressing MDCK I monolayers. **(A)** No CLDN15 expression was detected without doxycycline treatment. Following doxycycline treatment, wildtype CLDN15^WT^ or CLDN15^D55E^ and CLDN^D55N^ mutant was expressed at the tight junctions between cells (scale bar = 20 µm). **(B)** CLDN15 expression significantly reduced TER (mean ± SEM; Student’s unpaired two-tailed *t* test: CLDN15^WT^, *n* = 4 technical replicates, representative of six separate experiments, *P* = 2.6 × 10⁻¹²; CLDN15^D55E^, *n* = 8 technical replicates, representative of two separate experiments = 5.1 × 10⁻⁸; CLDN15^D55N^, *n* = 4 technical replicates, representative of two separate experiments, *P* = 2.1 × 10⁻⁸) **(C)** and increased relative permeability of sodium to chloride (PNa^+^/PCl^-^) (mean ± SEM; Student’s unpaired two-tailed *t* test: CLDN15^WT^, *n* = 4 technical replicates, representative of six separate experiments, *P* = 3.6 × 10⁻^7^; CLDN15^D55E^, *n* = 8 technical replicates, representative of two separate experiments, *P* = 0.0018; CLDN15^D55N^, *n* = 4 technical replicates, representative of two separate experiments, *P* = 4.2 × 10⁻^6^).

To assess the contribution of CLDN15 DFF55 to channel size selectivity, we measured biionic potentials after basolateral substitutions of different sized cations for Na^+^. This allowed calculation of the permeability of the substituted cation relative to Cl^-^. The data are shown normalized to Cl^-^ (Fig. 2A) or Na^+^ (Fig. 2B). In all cell lines studied, without induced CLDN15 expression, permeabilities of different sized cations were low and size non-selective, consistent with the cell lines have no confounding selective conductance pathway for ions studied. For CLDN15^WT^ expressing monolayers, peak permeability was for cations with radii ∼1.5Å (K^+^, Rb^+^, and Cs^+^) and was minimum for ions greater than radius 2.5Å (TMA^+^ and TEA^+^). CLDN15^D55E^ expressing monolayers exhibited a selectivity profile with permeability decreasing as radii increased (from Na^+^ to K^+^, Rb^+^, and Cs^+^), with Cs^+^ permeability being about half of peak ion permeability. CLDN15^D55N^ expressing monolayers had a permeability profile that peaked at slightly higher radii than CLDN15^D55E^ (e.g. close to K^+^). The most dramatic effect of CLDN15^D55E^ mutation is an overall depressed cation permeability relative to Cl^-^ permeability.

**Figure 2:**
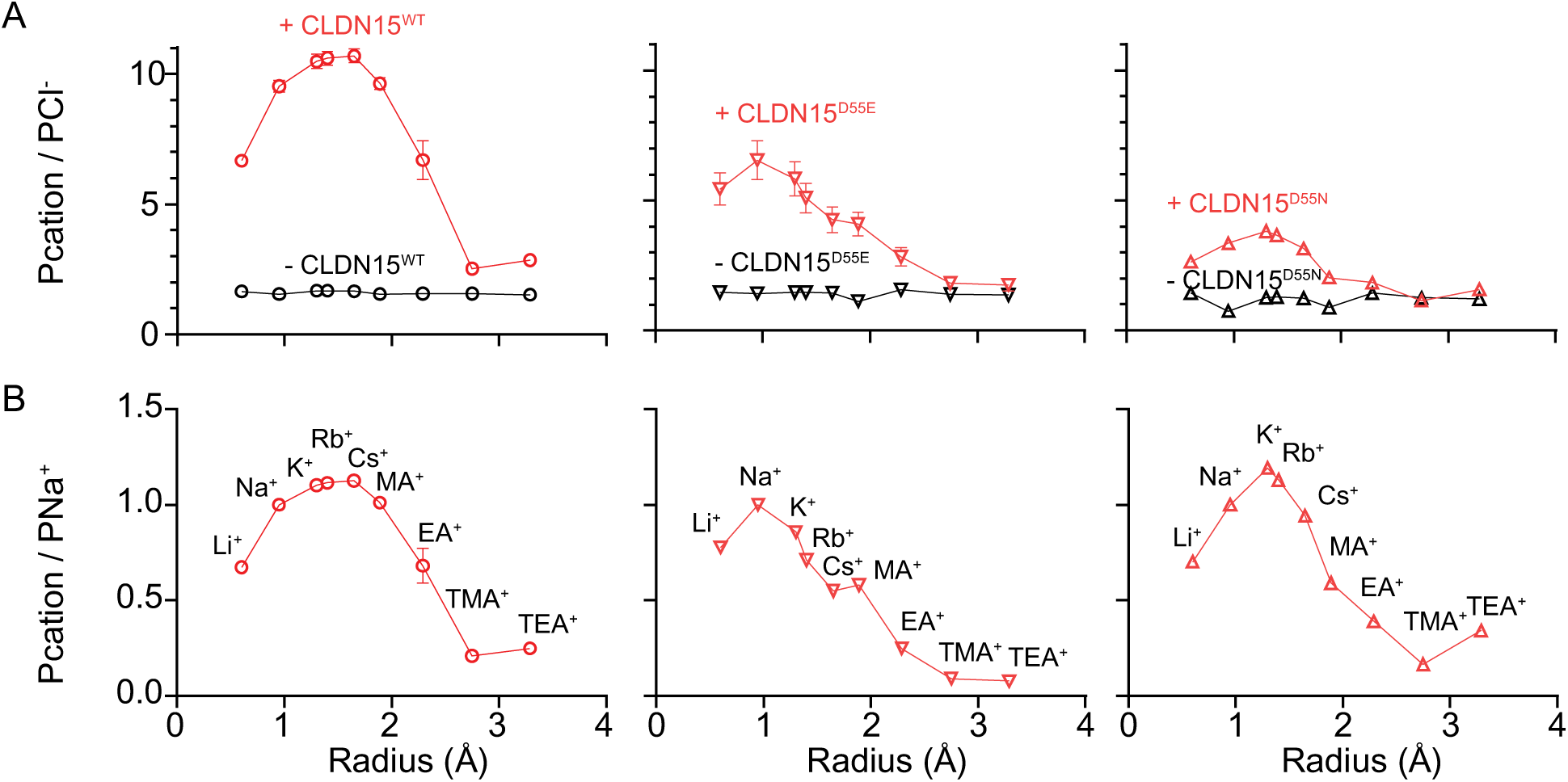
CLDN15^D55E^ and CLDN15^D55N^ mutants differentially affect pore charge and size selectivity profile. **(A)** Cation size selectivity was determined by using biionic potential measurements. CLDN15 expressing (red) and non-expressing (black) monolayers are shown. Cation permeabilities are normalized to chloride permeability to show charge selectivity as a function of cation size as well as the effects of D55E and D55N mutations. There was no charge selective barrier in control monolayers that did not express CLDN15, consistent with negligible baseline charge selectivity of the tight junction in parental MDCK I (mean ± SEM, n = 4) **(B)** The claudin-dependent size-selectivity profile for the same traces shown in A was calculated as the difference between cation permeability in monolayers that expressed or did not express claudin-15. Data are shown normalized to Na^+^ and illustrate significant differences in the shape of the curves between WT and mutant CLDN15 constructs. (mean ± SEM; CLDN15^WT^, *n* = 4 technical replicates, representative of six separate experiments; CLDN15^D55N^, *n* = 8 technical replicates, representative of two separate experiments; CLDN15^D55E^, *n* = 4 technical replicates, representative of two separate experiments).

### Molecular dynamic simulations allow us to study CLDN15 ion transport

We hypothesized that these differences in permeability properties between CLDN15^WT^, CLDN15^D55E^, and CLDN15^D55N^ are due to unique interactions of permeating ions within the selectivity filter at residue 55 and ion hydration. To test this, we turned to *in silico* modeling.

Previously, we have established and validated an all-atom molecular dynamic model of CLDN15 (Fig. 3A) that allowed us to simulate ion passage through the CLDN15 channel (Samanta et al., 2018).

**Figure 3:**
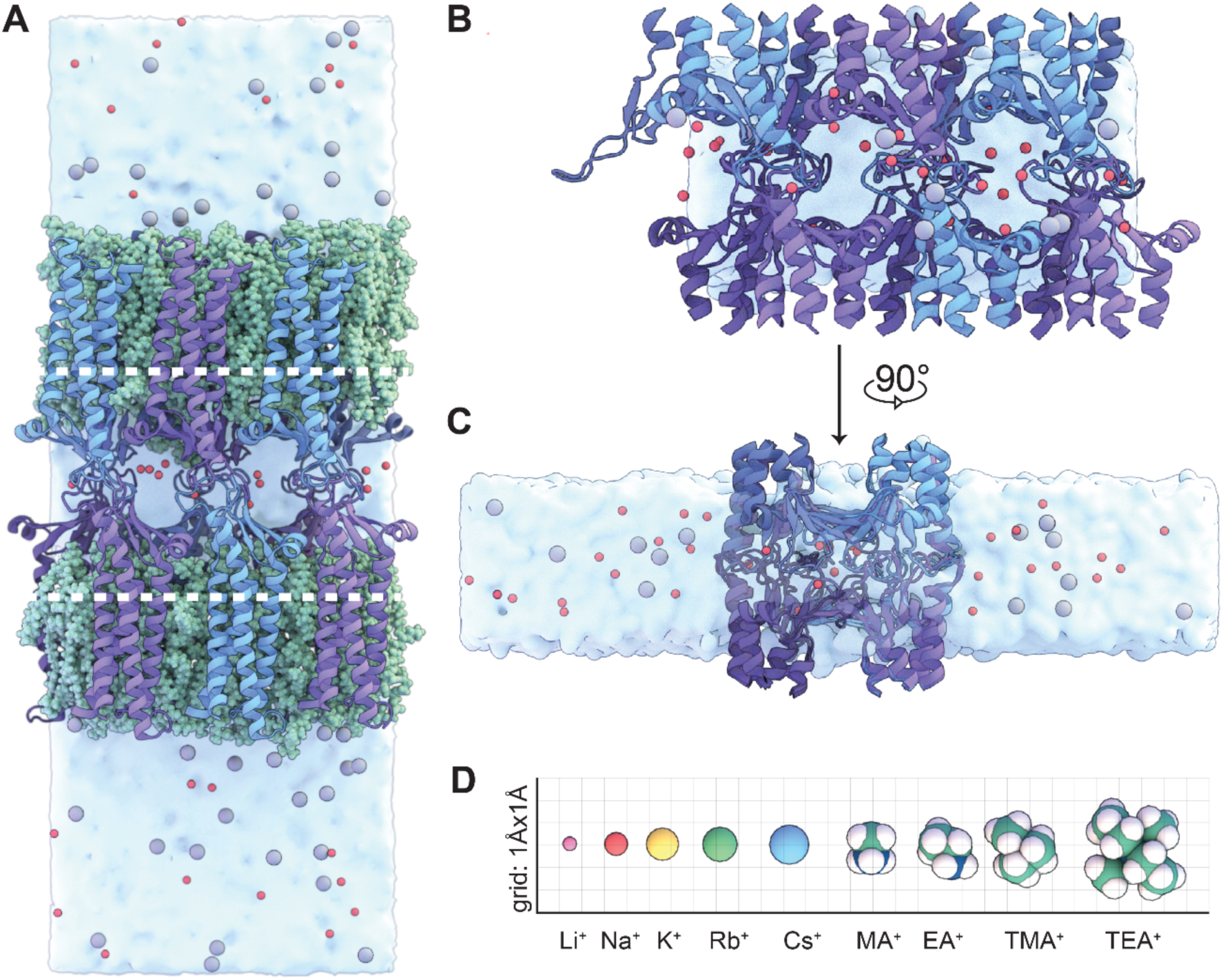
The reduced CLDN15 model allows for analysis of pore permeability properties. **(A)** A snapshot of the full CLDN15 model. The model contains two parallel lipid bilayers (green) and three CLDN15 pores in the unit cell. Na^+^ and Cl^-^ ions are represented in red and gray respectively. The white dotted lines indicate the section of protein removed in preparation for the reduced model. **(B)** View of the channels in the reduced CLDN15 model, in which the transmembrane helices and two lipid bilayers have been entirely removed. There are ∼15 Å of vacuum above and below the water slab and the water molecules in the slab are constrained to their initial volume, preventing them from diffusing into the vacated membrane region. This model includes the ion conduction pores, surrounding solutions, and part of the transmembrane helices. **(C)** Side view of the reduced system model after a 90° counterclockwise rotation. The solutions on the two sides represent apical and basolateral solutions, however, it must be noted that the channel is symmetric along the pore. **(D)** Cations used in comparative simulations, plotted by size in Angstroms.

However, this model is unsuitable for our intended modeling due to significantly increased computational need, which prevents us from performing multiple long simulation runs with reasonable time resolution.

To address this challenge, we attempted to create a reduced molecular dynamics model of CLDN15 ion conduction pore, aiming at increasing simulation throughput while preserving model fidelity, which would allow us to perform longer simulations at reduced computational cost, while maintaining simulation accuracy.

We created a reduced model that retains all the amino acid residues within the pore region and the surrounding environment but omits the transmembrane regions that anchor the protein within the lipid bilayer (Fig. 3B,C). Through this reduced model we achieved a six-fold improvement in computational efficiency compared to the full model. This substantial increased throughput would permit us to study conductivity of a range of monovalent cations (Fig. 3D) for all three CLDN15 forms studied *in vitro*.

We first compared structural and functional properties of the full model of CLDN15 (Fig. 3A to the reduced model (Fig. 3B,C). Similar to the full model, in the reduced model, each channel is composed of eight protomers: β-sheets from four primary protomers to form opposing (top and bottom) walls of the pore, and ECS2 (residues 149–150) and β1β2 loops (residues 34–42) within ECS1 (39–42) from four auxiliary protomers. The structural components of these protomers effectively seal the space between these channels. The stabilities and secondary structures of the models were comparable. For the reduced model, the time-averaged backbone RMSD relative to its initial structure was 1.8±0.05 Å over 105 ns, which is comparable to the average RMSD (1.57±0.16 Å) of the full model RMSD relative to its initial structure. Over the last 14 ns of equilibration, time-averaged per-residue RMSD and RMSF values remain reasonably low, with expected peaks for extracellular residues (Fig. 4A,B). Structurally, the organization of CLDN15 pores were identical in both models.

**Figure 4:**
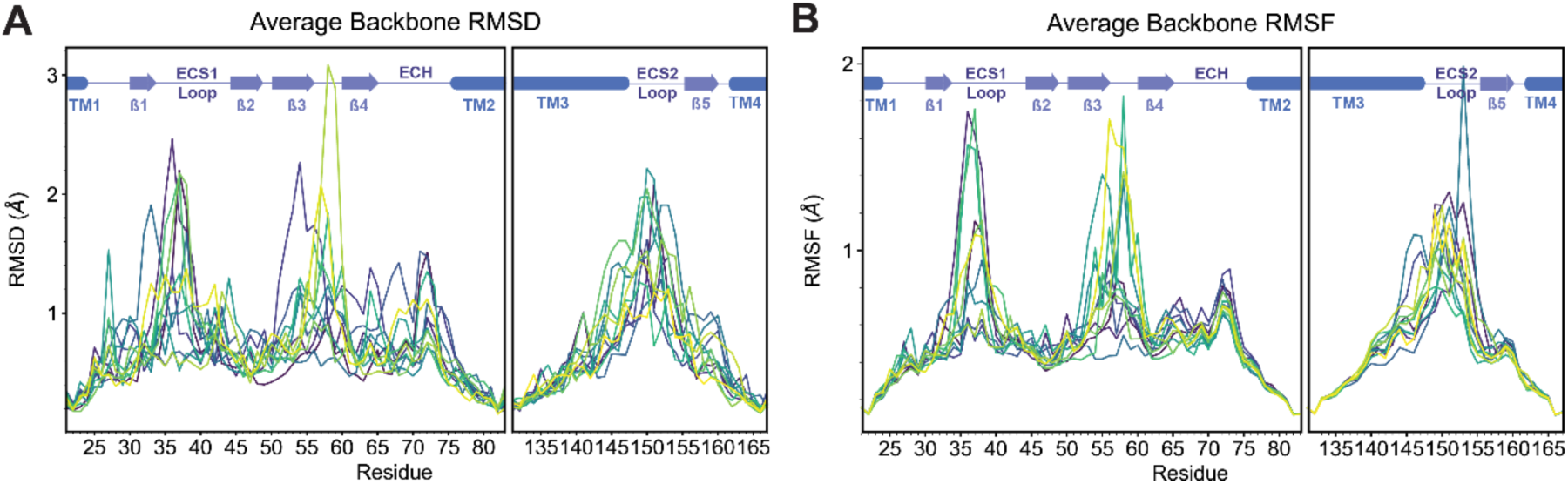
Structural stability of the equilibrated CLDN15 reduced model: **(A)** Average backbone RMSD from the last 14 ns of the CLDN15 reduced model 105 ns equilibration, colored by each of 24 monomers. **(B)** Average backbone RMSF from the last 14 ns of the CLDN15 reduced model 105 ns equilibration, for each of 24 monomers.

We next compared cation conductance in the CLDN15 pore through simulations of ion transport in the presence of a voltage gradient (Gumbart et al., 2012). In both models, the ion conduction pore was fully hydrated with ions freely entering and leaving the pore, and no ions leaking perpendicular to the pore were observed in simulations, judged by visual inspection of simulation trajectories. Simulations using our full model and reduced models showed similar single-pore conductance of Na^+^ (55.8 ± 16.8 pS in full model, and 56.7 ± 8.8 pS in reduced model).

To more comprehensively compare the performance of both models, we investigated the relationship between cation size and conductivity by examining the current-voltage relationship for cations of different sizes. This analysis allowed us to determine the conductivity profiles as a function of ion radius for both the full model (Fig. 5A, C) and the reduced model (Fig. 5B, D). Peak conductance values occurred at a cation radius of approximately 1.5 Å (K^+^, Cs^+^, and Rb^+^) in both models, which corresponds well with peaks in size-selective permeability determined in CLDN15^WT^ expressing monolayers.

**Figure 5:**
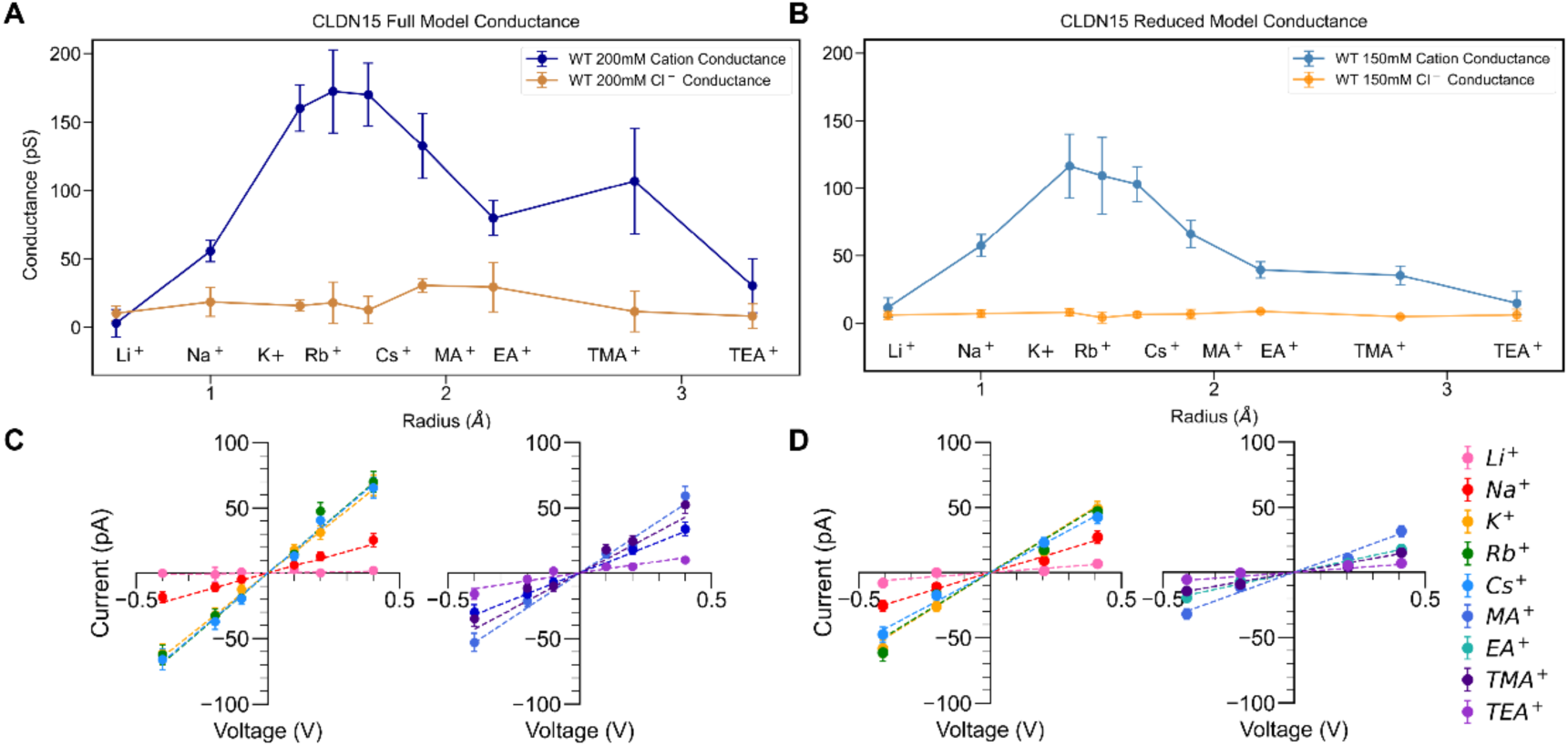
CLDN15 permeability profile is comparable between full and reduced CLDN15 models. **(A)** Single-channel cation and anion conductance of CLDN15^WT^ full model. **(B)** Single-channel cation and anion conductance of CLDN15^WT^ reduced model Replicas 1 and 2. **(C)** I-V plots of the full model CLDN15^WT^ for ±0.1, ±0.2, and ±0.4V. **(D)** I-V plots of the reduced model CLDN15^WT^ at ±0.2V and ±0.4V.

Furthermore, similar to cell culture data, Cl^-^ conductance was much lower than cation conductance in both the full model and reduced CLDN15^WT^ models. CLDN15 ^WT^ permeability profiles follow the pattern P_K_ ≈ P_Rb_ ≈ P_Cs_ > P_Na_ > P_Li_ for both full and reduced *in silico* models (Fig. 5A,B), consistent with biionic potential measurements (Fig. 2A). These results are consistent with the permeability profiles observed in MDCK II monolayers, where the major claudins were knocked out (claudin quintuple KO) and claudin-15 was re-expressed (Pouyiourou et al., 2025).

Taken together, our analyses showed the reduced and full model have similar CLDN15 pore organization, structural stability, and permeability of different sized ions, with significantly decreased model complexity. This newly developed reduced CLDN15 model enables us to effectively investigate structural and functional requirements for the CLDN15 permeability filter to control pore function.

### D55 Mutations Alter Charge and Size Selectivity of CLDN15 in Silico

Our cell culture data (Fig. 2) revealed differences in the size selectivity of conductance when D55 was mutated to E or N. We hypothesized that these differences were attributable to unique pore interactions between permeating cations and the selectivity filter as well as differences in the size of the pore. To evaluate the mechanism behind these differences, we created reduced CLDN15^D55E^ and CLDN15^D55N^ models and compared their simulated cation conductance profiles to those of CLDN15^WT^ (Fig. 6A).

**Figure 6:**
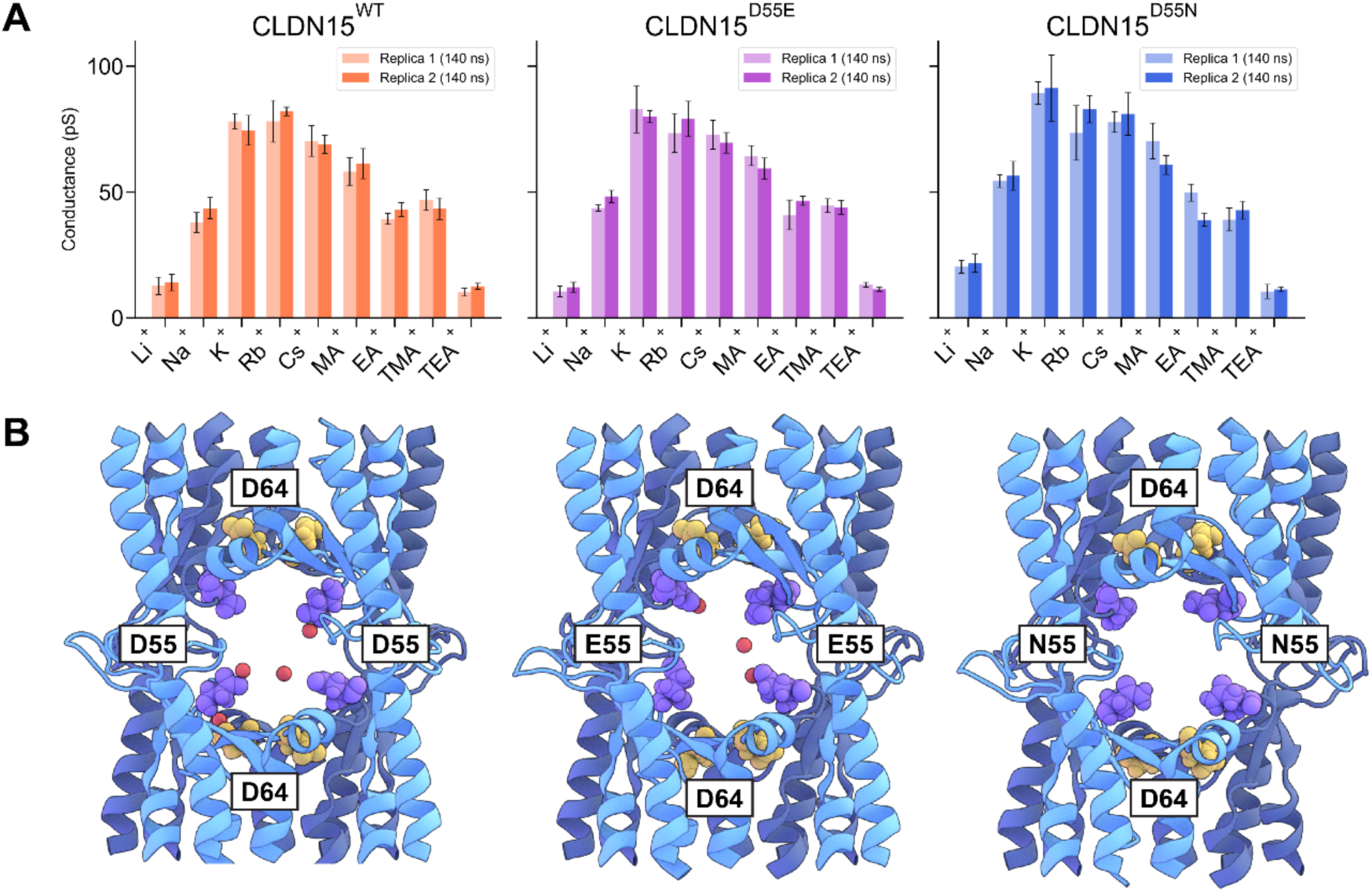
Mutant conductivity. **(A)** Single-channel cation conductance profiles calculated from 140 ns constant voltage simulations (V = ± 800, 1200 mV) from the reduced model CLDN15^WT^, CLDN15^D55E^, and CLDN15^D55N^ systems at 100 mM cation concentration. Two replicas were simulated to reduce statistical variation. **(B)** Snapshots of CLDN15 WT, D55E, and D55N channels taken from simulations. D64 (yellow), D55 (purple), and Na^+^ (red) are shown.

In CLDN15^WT^ simulations, we observed peak cation conductance at the size of Rb^+^ (1.5 Å in radius) (Fig. 6A). The CLDN15^WT^ pore features a cage-like structure comprising four D55 residues (Fig. 6B), which binds to and regulates the passage of ions to define permeability(Alberini et al., 2018; Samanta et al., 2018). Four D64 residues are represented in Fig. 6B to show the difference between pore-lining (D64) and pore-facing (D55) residues. CLDN15 mutants studied also adopted identical pore organization (Fig. 6B). CLDN15^D55E^ mutation led to a slight leftward shift in peak conductance, with decreasing conductance for cations larger than K^+^ and Rb^+^ (radius of 1.4 – 1.5 Å). This general permeability pattern was also observed in our cell culture studies (Fig. 2). In our simulations, it is challenging to appreciate slight differences in peak cation conductance between CLDN15^D55E^ and CLDN15^D55N^, which we observed *in vitro*, due to statistical fluctuations and the finite length of the simulations. These data illustrate the importance of charge at residue 55 on CLDN15 permeability.

### Pore Radius Differences Are Not a Major Determinant of CLDN15 Selectivity

We hypothesized that the slight leftward shift in the peak Na^+^ permeability of simulated permeability profiles of CLDN15^D55E^ relative to CLDN15^WT^ could be due to a narrower pore radius for CLDN15^D55E^ in comparison to CLDN15^WT^ or CLDN15^D55N^ (Fig. 7). To assess this, we calculated the average pore radius profile for three channels along its conductivity pathway. We compared the average pore radii over 140 ns trajectories of CLDN15^WT^, CLDN15^D55E^, and CLDN15^D55N^, which were 3.80 ± 0.51, 3.88 ± 0.67, and 3.87 ± 0.62 Å, respectively, with an average minimum pore radii of 2.18 ± 0.50, 2.18 ± 0.50, and 2.17 ± 0.59 Å. Thus, we observed no significant difference in average pore radius or minimum pore radius between the three systems.

**Figure 7:**
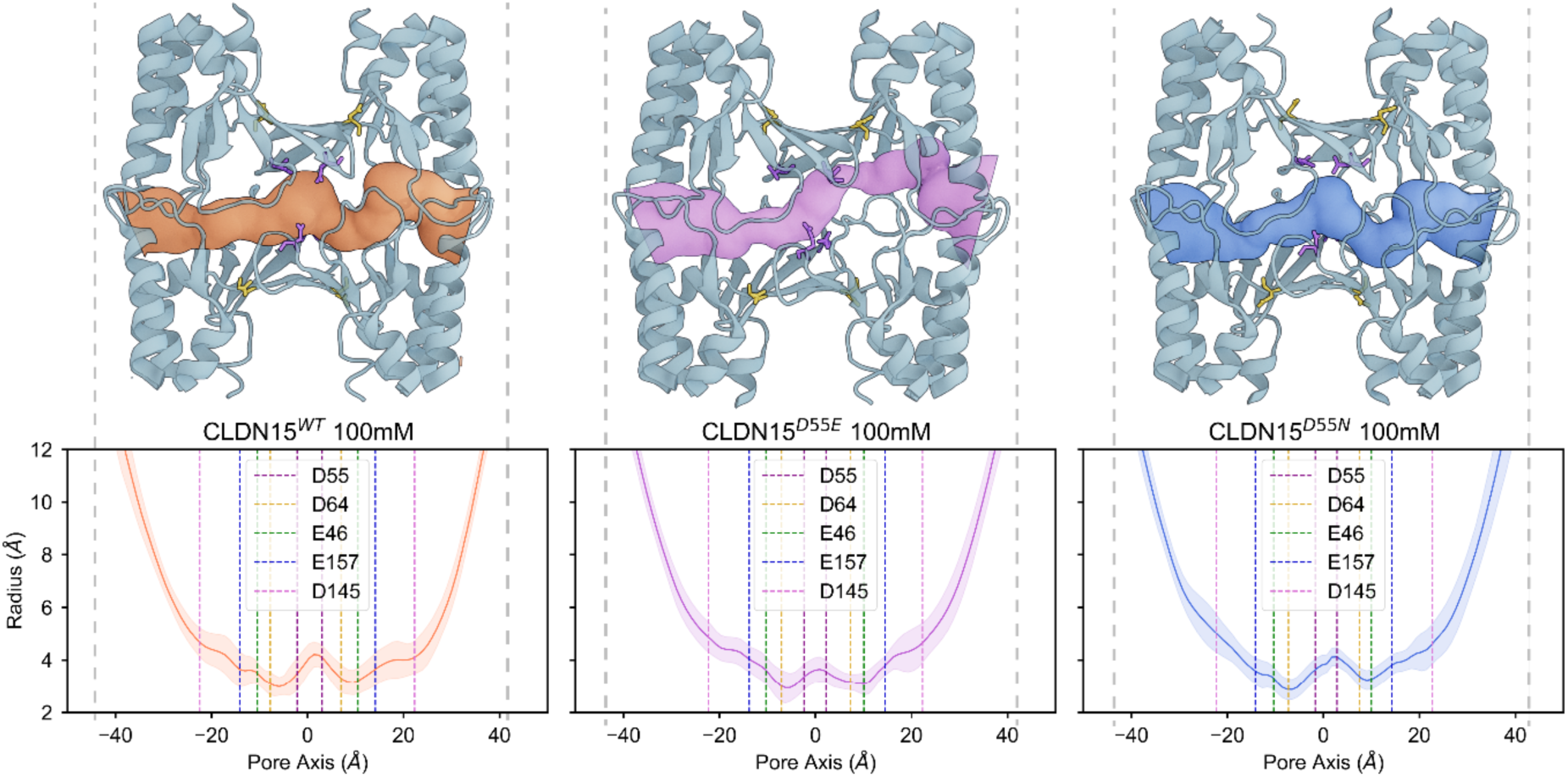
Pore radius. Snapshots include the primary tetramer (pale blue) and pore surface generated from HOLE2 analysis for CLDN15^WT^, CLDN15^D55E^, and CLDN15^D55N^ colored orange, purple, and blue, respectively. The binding site at residue 55 (purple) and D64 (yellow) is shown in Licorice representation without hydrogen atoms for each snapshot. Average and standard deviation for pore radius over 140 ns of voltage (−800 mV) simulation for CLDN15^WT^ (orange), CLDN15^D55E^(purple), and CLDN15^D55N^ (blue). The average Cα positions of negatively charged pore-lining residues D55, D64, E46, E157, and D145 are represented by dotted lines in the pore radius plots.

### D55 plays a key role in defining cation contact time with pore-lining residues

The simulations above suggest that pore size differences among the three CLDN15 models are negligible in *in silico* models. We hypothesized that differences in permeability among CLDN15^WT^, CLDN15^D55E^, and CLDN15^D55N^ are largely dominated by interactions of cations with negatively charged pore residues. To test this possibility, we determined the contact time of different amino acids to cations, defined as the number of cations bound to a specific residue at any given moment in each of the three CLDN15 models. For Na^+^ (Fig. 8A), the negatively charged residues D55 in CLDN15^WT^ and E55 in CLDN^D55E^ exhibited strong interactions with Na^+^, with an average of two sodium ions bound to D55 or E55 at all times (Fig. 8A,B). Fig 8B shows an overall pore occupancy of approximately five sodium ions. In contrast, overall pore occupancy for CLDN15^D55N^ was depleted to two cations per channel (Fig. 8B), which could effectively reduce the capacity of CLDN15 channel to conduct cations. Notable pore-facing interaction sites include E157, N61, S56, D55, and I39 (Fig. 8C). Contact time plots (Fig. 8A) show significant interaction between cations and the pore-lining residue D64 in both CLDN15^WT^ and CLDN15^D55E^ models, although D64 interactions are less pronounced in CLDN ^D55E^ models. These plots also show that hydrophilic residue N61 increased contact time in CLDN15^D55E^ channels (from 36% in CLDN15^WT^ to 76% in CLDN15^D55E^) (Fig. 8A). Additional secondary interactions were found with residues from the 𝛽1/𝛽2 loop (residues 35-42) including ECS1, with hydrophobic amino acids I39, T40, and V38 each binding Na^+^ approximately 30% of the time. The negatively charged E157 residue, positioned at the channel opening, was bound by Na^+^ ∼40% of the time (Table 2).

**Figure 8:**
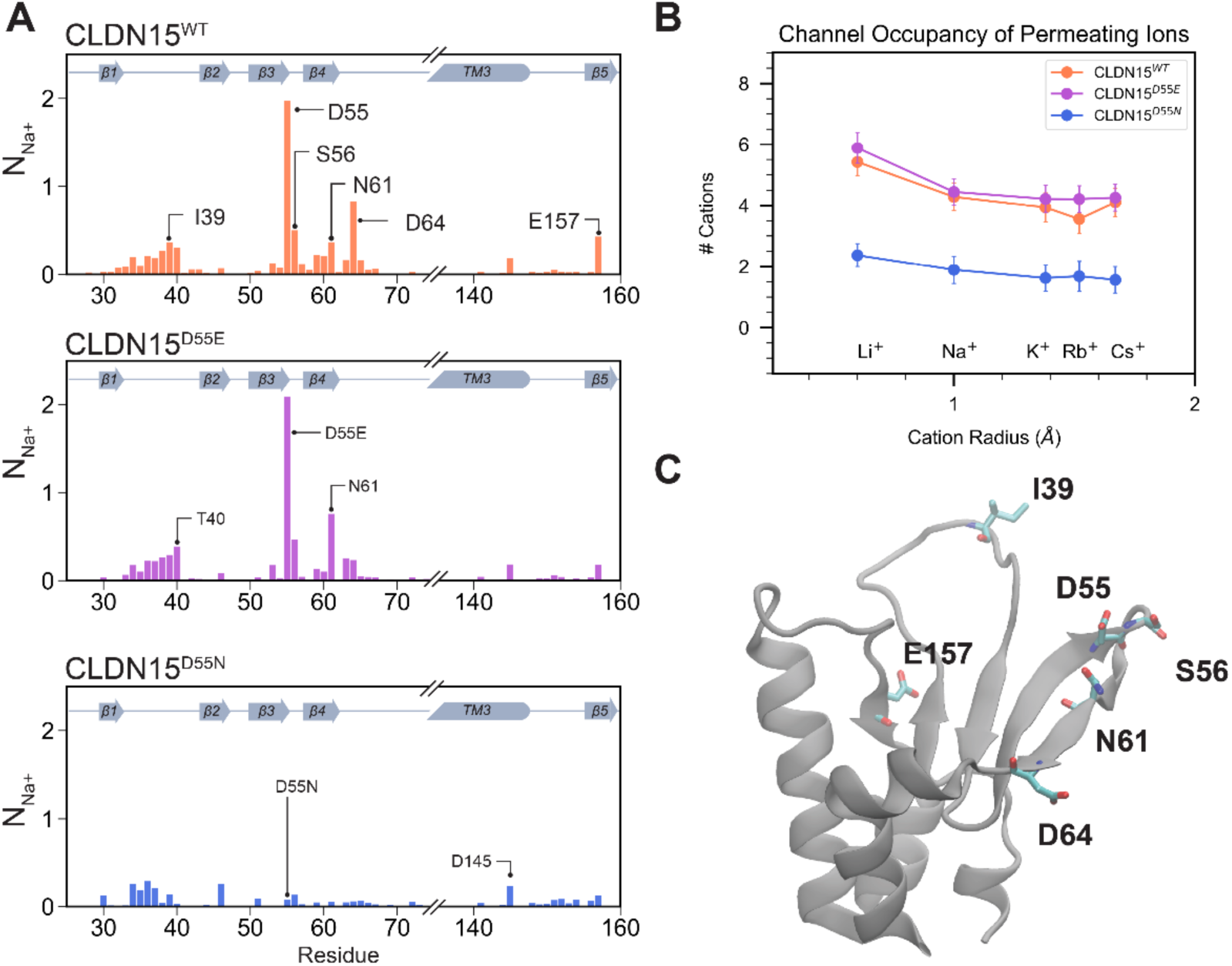
CLDN15 D55 is the principal cation binding site, and this is abrogated in the D55N mutation. **(A)** Plots show contact time data as the average number of Na^+^ ions that are found within 4.0 Å of CLDN15 amino acid residues in voltage simulations (V = ± 800, 1200 mV) for CLDN15^WT^, CLDN15^D55E^, and CLDN15^D55N^. **(B)** Average number of cations (Li^+^, Na^+^, K^+^, Rb^+^, Cs^+^) within each pore for CLDN15^WT^, CLDN15^D55E^, CLDN15^D55N^, calculated from voltage trajectories. **(C)** Position of key cation interacting sites on CLDN15: I39, D55, S56, N61, D64, E157.

**Table 2.** Average number (N) of Na^+^ ions bound to pore-lining residues within CLDN15^WT^.

| Residue | $N \pm SD$ | Residue | $N \pm SD$ | Residue | $N \pm SD$ |
| --- | --- | --- | --- | --- | --- |
| D55 | $1.97 \pm 0.16$ | E157 | $0.43 \pm 0.25$ | T40 | $0.30 \pm 0.15$ |
| D64 | $0.83 \pm 0.33$ | N61 | $0.36 \pm 0.12$ | V38 | $0.27 \pm 0.12$ |
| S56 | $0.50 \pm 0.2$ | I39 | $0.36 \pm 0.11$ | V59 | $0.21 \pm 0.12$ |

### Dehydration of Alkali Cations Induced by D55 Aspartic Acid Residues: A Critical Step for Selectivity During Permeation Through the CLDN15 Pore

Our findings show the critical role of D55 in ion conductance and selectivity within the CLDN15 pore. To further investigate the mechanisms by which D55 influences cation selectivity, we focused on the process of dehydration—a key step in ion permeation that directly impacts ion-channel interactions.

We subsequently analyzed how alkali cations lose their hydration shells while traversing the CLDN15 pore and quantified the contributions of pore-lining residues to this process. In addition to calculating the number of coordinating water molecules, we determined the number of oxygen atoms from CLDN15 that interact with cations. We observed that regardless of ion size, all cations are partially dehydrated by the channel as they traverse the pore. For all alkali cations to permeate, they must lose at least 2 coordinating water molecules to pass through the channel. Smaller cations such as Li^+^, which has one of the lowest conductivities, are in close proximity of the binding site(s) and remain bound to the channel as they permeate through the pore. On the other hand, Rb^+^, which has one of the highest conductivities, faces the smallest dehydration penalty with respect to the size of its hydration shell (Fig. 9). Larger ions such as Rb^+^ bind readily to the D55(or E55)/N61/D64 binding site(s), but the size of its hydration shell permits hopping between the binding sites along the pore axis and pass through the pore more readily than smaller cations. In the absence of the negatively charged D55 (or E55) residue, in the D55N mutant, permeating cations maintain their contact with pore lining residues N61 and D64 and pass through the pore unaffected (Fig. 9C). The lower cation selectivity of the D55N mutant is mostly due to higher Cl^-^conductivity as shown previously (Samanta et al., 2018)).

**Figure 9:**
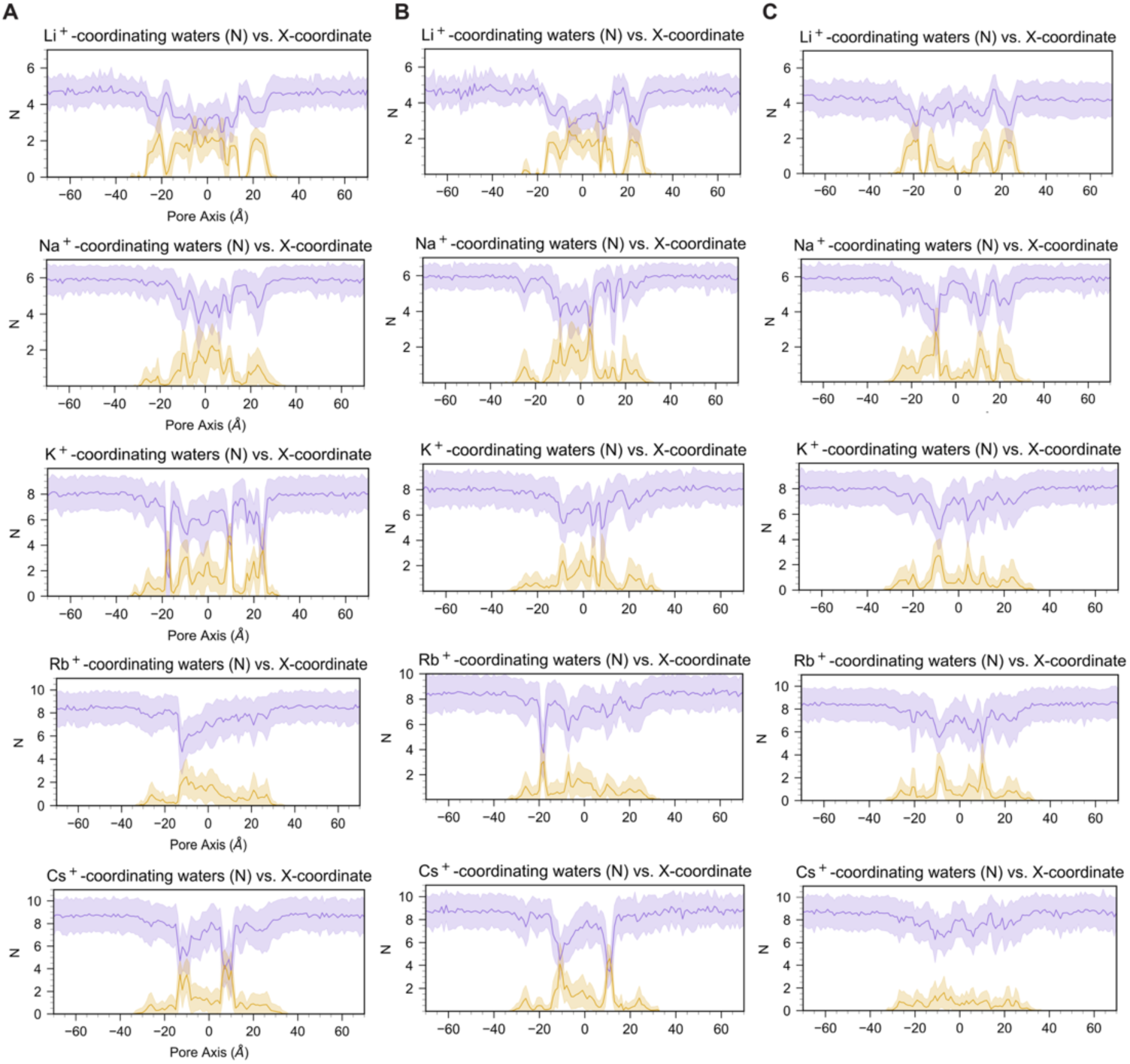
Dehydration of permeating alkali cations for CLDN15. 40 ns trajectory analysis of voltage (−1.2 V) simulations for **(A)** CLDN15^WT^ 100 mM X^+^, **(B)** CLDN15^D55E^ 100 mM X^+^, and **(C)** CLDN15^D55N^ 100 mM X^+^ (where X^+^ is alkali cation Li^+^, Na^+^, K^+^, Rb^+^, or Cs^+^). The average and standard deviation of coordinating water molecules in the first hydration shell (purple) and oxygen atoms from protein residues (gold) across the pore axis, averaged over 3 channels in the unit cell.

Partial dehydration by the binding site and transient interactions with specific pore-lining residues contribute to the cation permeation process through CLDN15 channels, highlighting the importance of understanding this complex network of interactions in regulating selectivity and permeability.

As the ionic radii of alkali cations increases, so does the number of coordinating waters (Fig. 10A). Dehydration events occur on a timescale of nanoseconds (Fig. 10B). Our analyses show that when Na^+^ entered the pore with ∼6 coordinating water molecules, it loses ∼2 coordinating water molecules as it passed through the selectivity filter formed by four aspartic acid residues (D55) in the center of the CLDN15 pore. D55 was the primary residue involved in these dehydration events. (Fig. 10C).

**Figure 10:**
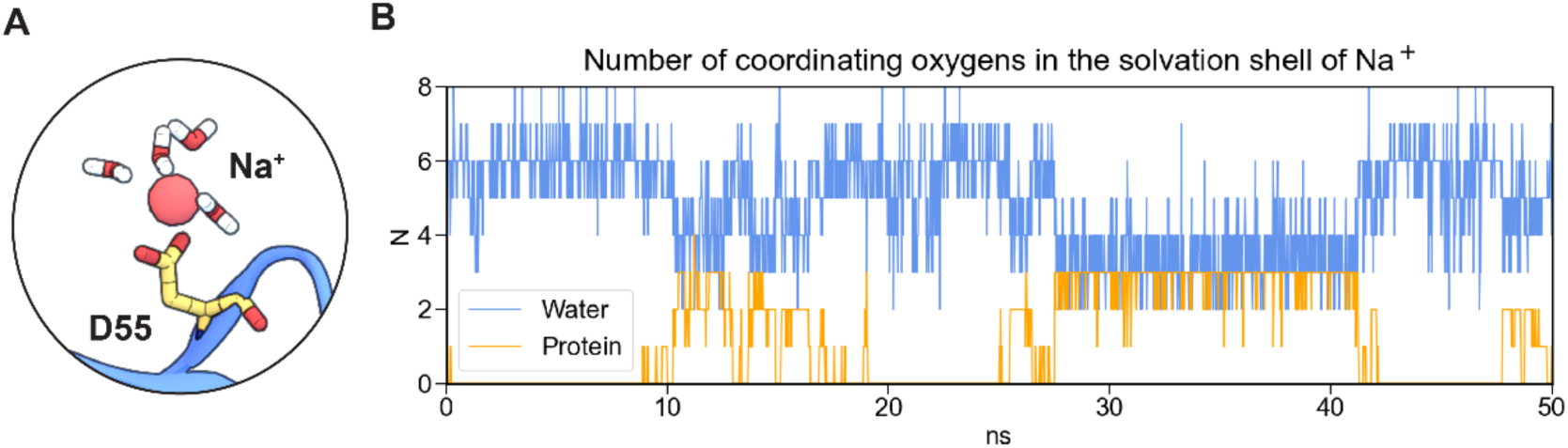
Dehydration of permeating cations along the pore. **(A)** Snapshot of dehydrated Na^+^ bound to D55, having 4 water molecules in its hydration shell. Two water molecules and their oxygen is replaced by two contributing oxygens from D55 sidechain. **(B)** The number of Na^+^ coordinating oxygens fluctuates over 50 ns trajectory as Na^+^ is dehydrated by pore-lining residues.

A detailed analysis of Na^+^ dehydration dynamics revealed fluctuations in the number of coordinating water molecules of Na^+^ as they are replaced by protein oxygen atoms, primarily from sidechains (Fig. 9A). At least one-third of partially dehydrated Na^+^ ions interacted with the D55 selectivity filter at any given time, with transient binding to other residues, including D64, N61, A53, E157, F65, T40, N37, S56, D145, S60, G36, I39, V38, and E46, occurring with decreasing frequency (Fig. 11A). The sidechain carboxyl group from negatively charged D55 and the sidechain oxygen of the polar N61 coordinate partially dehydrated Na^+^ complexes at the D55 binding site (Fig. 11B). T40 on ECS1 is near the D55 binding site and similarly stabilizes the Na^+^ complex via its backbone carbonyl oxygen (Fig. 11C). After passing through the selectivity filter, Na^+^ regains its full hydration shell as it exits the pore. However, the polar N37 and T40 are not located on the same monomers as the D55 binding sites.

**Figure 11:**
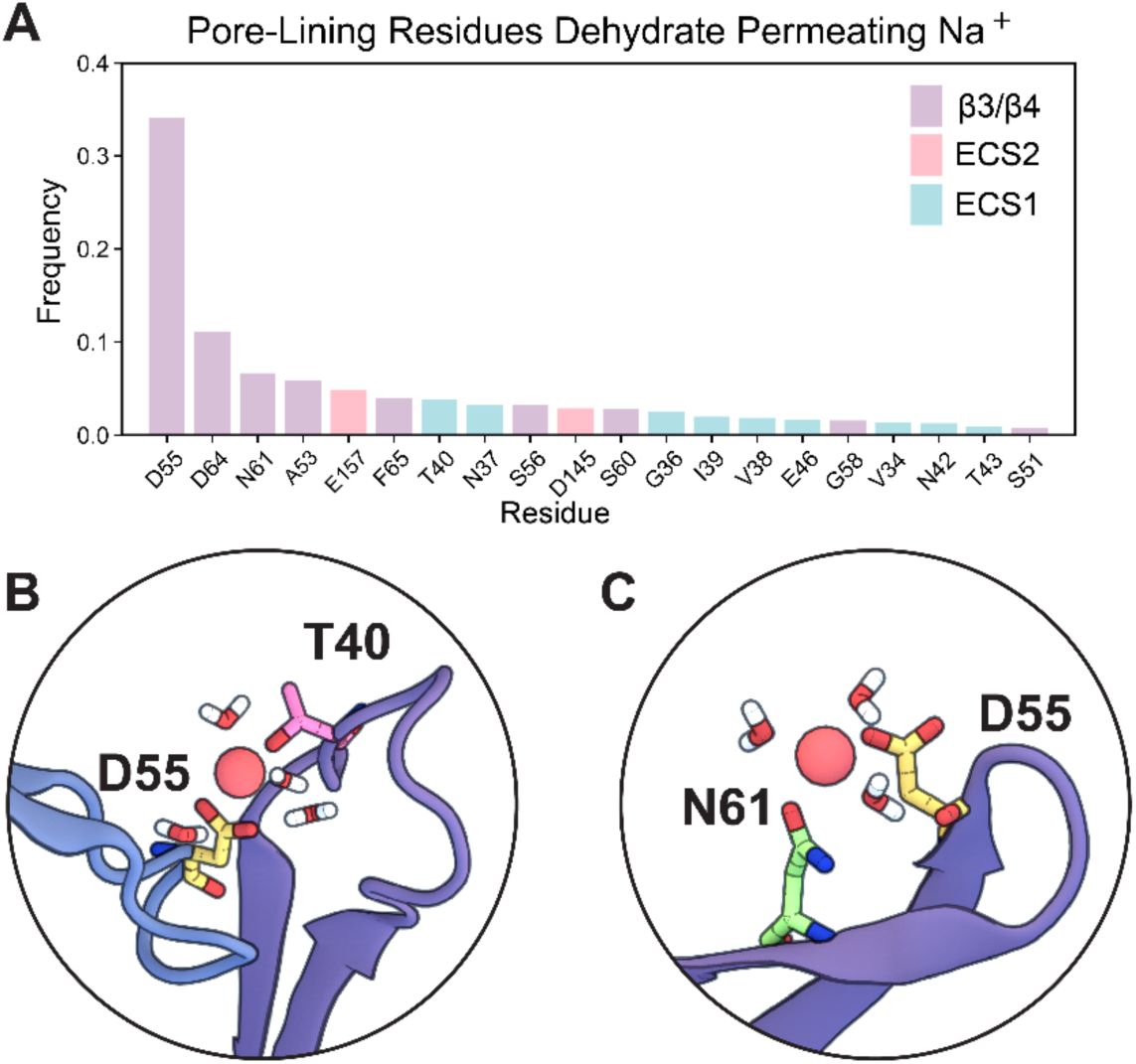
Pore-lining residues dehydrate permeating Na^+^. **(A)** Frequency of pore-lining residues which assist in Na^+^ dehydration as it permeates the channel. **(B)** Snapshot of neighboring residues D55 and N61 and partially dehydrated Na^+^ with five coordinating water molecules. **(C)** Snapshot of residue D55 on one monomer and T40 from the ECS1 of an adjacent monomer stabilizing a dehydrated Na^+^ with three coordinating water molecules.

The octameric arrangement of subunits and relative positions of pore-lining residues highlights the importance of dehydration interactions coordinated by multiple pore-lining residues. This arrangement, defined in the context of the CLDN15 pore model (Fig. 12A), examines the contribution of 8 monomeric subunits to a single pore. The D55 binding sites are formed by four central monomers (A1, A2, A3, A4) (Fig. 12B) arranged in a tetramer (Fig. 12C). Overlapping tetramers create a hydrophobic seal on either side of the pore. ECS1 from four laterally adjacent monomers (L1, L2, L3, L4) features alternating hydrophobic pore-lining sidechains (I39, V38) and polar uncharged sidechains (T40, N37, G36) that line the mid-pore region (Fig. 12B-C). We observed a constriction of the pore flanking either side of the D55 binding sites, lined by hydrophobic sidechains V38 and I39 (ECS1) (Fig. 12B). Similarly, the pore openings are flanked by hydrophobic ECS2 residues P149, L150, and Y151 (Fig. 12B). This complex arrangement of the CLDN15 channel coordinates the electrostatic environment of the channel to permit cation diffusion.

**Figure 12:**
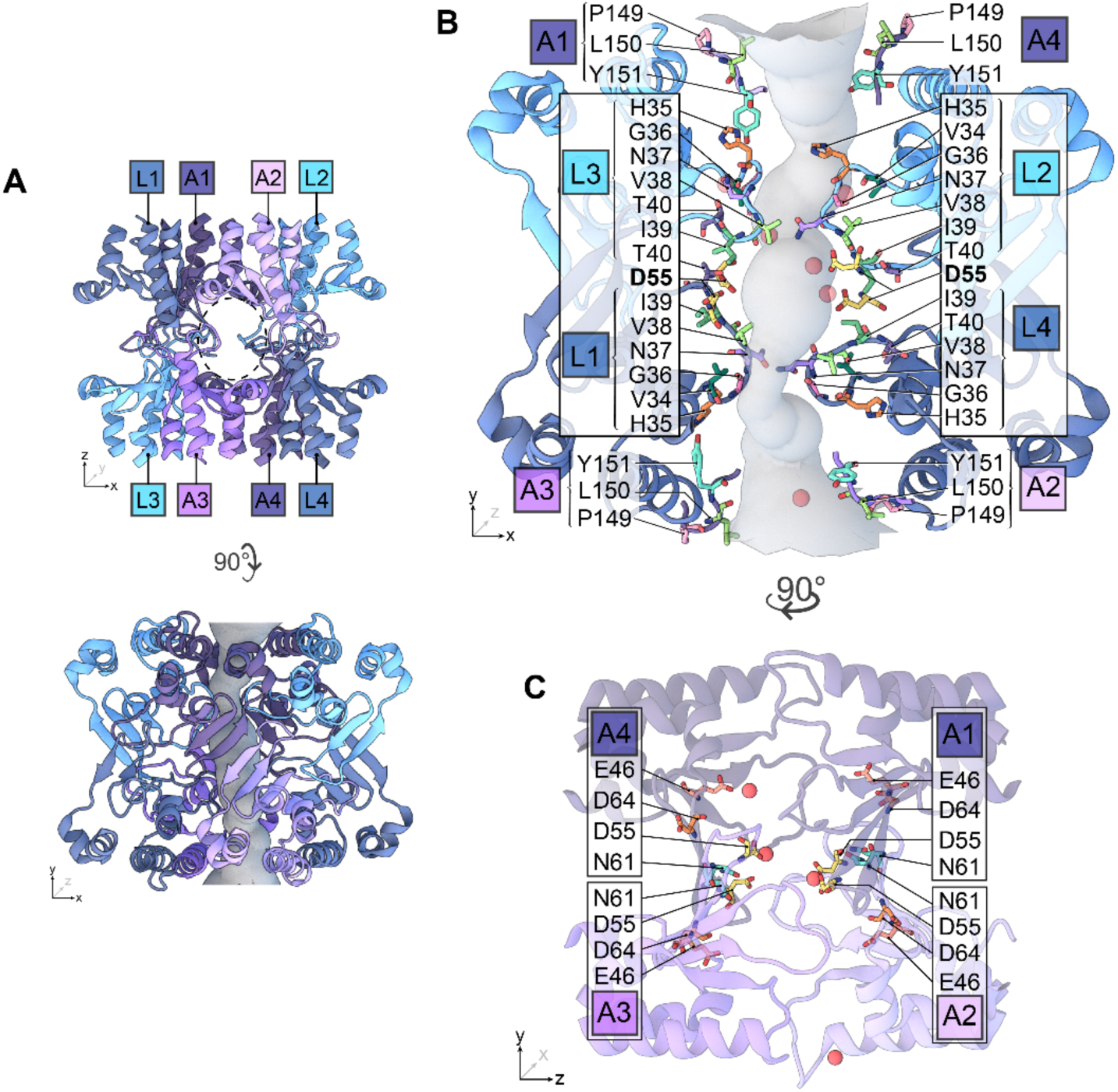
Pore-lining residues suggest octameric arrangement of CLDN15 monomers. **(A)** Octameric channel structure reveals contribution of lateral monomers (L1, L2, L3, L4) to central monomers (A1, A2, A3, A4). **(B)** Pore-lining residues include H35, G36, N37, V38, I39, T40 from β1β2 regions on lateral subunits as well as residues P149, L150, Y151 from ECS2 regions on primary subunits. **(C)** Sequence of negatively charged E46, D64, D55, and polar residue N61 located on the channel’s primary subunits.

## DISCUSSION

CLDN15 plays critical roles in nutrient, electrolyte, and water transport (Tamura, 2011; Tamura et al., 2008; Tsukita et al., 2019). It is well recognized that CLND15 creates a size- and charge-selective channel(Gonschior et al., 2022). Our previous study using all atom molecular dynamic simulations and cell culture studies demonstrated that the D55 amino acid residue forms the CLDN15 selectivity filter and serves as the primary binding site for cations. Within the CLDN15 channel, the four D55 amino acid residue side chains extend toward the center of the channel forming a cage-like structure that attracts cations. Despite these findings, the biophysical determinants of the D55 selectivity filter and how it may impact cation permeation remain poorly understood. We addressed this question by using a combination of cell culture and molecular dynamics simulation studies. Our investigation revealed key insights into these mechanisms.

To investigate how D55 creates the permeability filter and controls cation transport, we examined the *in vitro* behavior of WT CLDN15 alongside CLDN15^D55E^ and CLDN15^D55N^ mutants, determining their in vitro permeability properties and modeled ion flux. Our studies showed that although similar expression was achieved in high resistance MDCK cells, these CLDN15 channels have different permeability to alkali cations relative to Cl^-^. CLDN15^WT^ channels have the highest relative permeability to cations, with the relative permeability of K^+^, Rb^+^, and Cs^+^ to Cl^-^ being similar. In contrast, the CLDN15^D55E^ mutant exhibited slightly reduced cation selectivity and reduced permeability of K^+^, Rb^+^, and Cs^+^ relative to Na^+^. The CLDN15^D55N^ channel exhibited significantly lowered cation selectivity, with K^+^ showing the highest relative permeability amongst the tested cations. Notably, the CLDN15^D55N^ channel still retained some cation selectivity, indicating that D55 is not the only cation binding site within the CLDN15 channel. This is consistent with our previous finding that other weaker cation binding sites exist within the CLDN15 channel. These findings indicate that even a small change of D55E could impact relative cation selectivity by affecting the ability of the permeability filter to bind to cations and dehydration penalty for entering the pore.

Building on these *in vitro* findings, we developed a reduced CLDN15 model that contains the ion conduction pore of the CLDN15 complex and surrounding solution, with the transmembrane domains removed. Despite these simplifications, the reduced CLDN15 model preserves all atoms within the conductivity pathway and exhibits conductance and selectivity properties comparable to the full model. This streamlined model enabled us to efficiently explore hydration dynamics and cation interactions while retaining essential pore features, making it a robust tool for investigating ion transport. In this model, each channel is composed of eight monomers: four primary monomers contributing β-sheets that form the primary channel structure, and four auxiliary monomers—contributing ECS2 (residues 149–150) and β1β2 loops (residues 34–42)—that effectively seal the space between channels. This model offers a practical solution for studying claudin channels at atomic resolution while overcoming the computational challenges posed by full molecular dynamic models. **Its computational efficiency allowed us to investigate a broader range of conditions, such as ionic concentrations and voltage variations, which were critical for understanding how hydration influences ion transport.**

Although general permeability profiles of alkali cations were similar to *in vitro* measurements, differences in alkali cation permeability among CLDN15 mutants were less pronounced. This discrepancy may arise because certain factors, such as lack of polarizability in the force fields used in these simulations and the finite sampling time. Incorporating these additional factors could improve modeling accuracy in detailed ion permeation studies.

As alkali cation dehydration within ion channels can impact ion permeability, we determined how CLDN15 channel may impact alkali cation hydration. Simulations showed that fully hydrated cations are too large to pass through the CLDN15 pore. To permeate, cations such as Na⁺ must undergo partial dehydration, losing approximately two water molecules from their hydration shell to reduce size. All other alkali cations (Li⁺, Rb⁺, K⁺, Cs⁺) must also shed at least two coordinating waters to pass through the pore. Size differences in cation hydration shells impact conductivity, with larger cations like Rb⁺, K⁺, and Cs⁺ exhibiting high conductivity through the channel.

This dehydration process represents a critical energetic hurdle that is mitigated by a fine balance of contributions from the electrostatic forces created by D55 and the stabilizing effects of adjacent residues, such as carboxyl and backbone oxygens from D64 and the sidechain oxygen from N61. The negatively charged D55 residues play a central role in this process, destabilizing the cation’s hydration shell to facilitate dehydration while also stabilizing the partially dehydrated cations as they pass through the pore.

Polar uncharged residues such as N37, T40, S56, S60, and N61 provide additional stabilization, forming transient electrostatic interactions that guide the dehydrated ions through the channel and maintain ion selectivity. These extracellular regions emphasize the cooperative nature of ion transport in the CLDN15 channel, highlighting that efficient dehydration and stabilization are not exclusively dependent on the charged D55 residue. This observation held true across CLDN15^WT^, CLDN15^D55E^, and CLDN15^D55N^, indicating that a negatively charged residue at D55 is not strictly required for cation dehydration and subsequent permeation. Instead, pore-lining residues from the ECS1 regions contribute hydrophilic interactions that complement the function of D55. Furthermore, these regions arise from an arrangement of overlapping tetramers, in which four central monomers each contribute a D55 binding site to the channel and four adjacent monomers contribute their ECS1 loop regions to seal the lateral walls of the pore. The importance of these interactions in coordinating selective ion transport emphasizes the reliability of an overlapping tetramer arrangement.

The D55N mutation provided key insights into this mechanism, as it showed reduced charge selectivity and less effective dehydration of cations. This increased hydration correlated with reduced permeability, demonstrating the energetic importance of the electrostatic environment at D55. Abolishing D55 as a binding site led to a marked reduction in ion interactions with the channel, as reflected by decreased channel occupancy, shorter ion contact-time, and reduced overall conductivity. Conversely, the D55E mutation preserved the charge properties of D55 but altered the steric properties, subtly shifting the ion selectivity profile, but with no significant change in pore size in our model. These findings illustrate the interplay between charge and size selectivity at D55 and underscore the cooperative roles of adjacent residues in fine-tuning pore function.

The stabilization of dehydrated cations by these regions provides critical new insights into the cooperative roles of extracellular domains in ion transport through claudin channels, emphasizing that dehydration is not exclusively dependent on charged residues within the pore but also on hydrophilic interactions from adjacent structures. Such insights into the hydration process are critical for understanding how claudin mutations or alterations in pore structure may lead to dysregulation of ion transport in disease states. Future studies could investigate whether therapeutic interventions targeting hydration dynamics, such as small molecules that mimic or enhance these interactions, can restore normal claudin function in pathological conditions.

As a whole, our study validates that our CLDN15 model delineates the pore-lining residues of claudin-15 reasonably well. However, the structure of the CLDN15 pore is yet to be determined experimentally and the model stands to be refined as new experimental data emerges. For instance, future work could integrate cryo-EM or crystallographic data to further validate and refine our computational predictions, bridging the gap between *in silico* and experimental approaches. One observed limitation of the model is that we were not able to detect the leftward shift in size selectivity in the mutants, which we had confidently observed *in vitro*. This was partly due to the size of the error bars, but it could also be caused by inaccuracies of the simulation forcefields. For example, non-polarizable forcefields used in our simulations, inherently carry some degree of error in representing ion, protein, and water interactions. The finite length of the simulation, especially for ions with low conductivities, plays an important role in simulation accuracy. The accuracy of the simulation data improves with increasing simulation time as the number of transport events (n), especially for the ions with low conductivity (error ∼ (1/√𝑛)) (Aksimentiev et al., 2004; Aksimentiev and Schulten, 2005; Khalili-Araghi et al., 2013). These limitations highlight the need for improved computational tools and methodologies, such polarizable force fields or enhanced sampling approaches, which could provide a more accurate representation of ion dynamics in tight junction channels.

In summary, our findings demonstrate that the amino acid residue D55 plays a pivotal role in defining the conductance properties of CLDN15. Beyond enhancing our understanding of the ion permeation characteristics of CLDN15, the reduced model we developed offers a valuable tool for high throughput screening of small molecules *in silico* for their ability to bind to CLDN15 channels and modulate their permeability. This approach can be extended to other claudin channels to investigate their transport properties, paving the way for the rational design of claudin mutants that selectively regulate the transport of ions and small molecules. In future studies, we will apply the same strategy to generate reduced models of other members of the claudin family, thus elucidating their channel structure and the molecular basis for their channel selectivity. These computational models can be used to develop claudin channel blockers, which may be beneficial therapeutically when it is desirable to limit paracellular water and ion transport. These insights hold promise for the development of targeted interventions that modulate claudin function, potentially leading to therapeutic strategies that improve barrier function and address diseases linked to tight junction dysfunction. The results and tools developed in this report will not only advance our basic understanding of the biophysical properties of CLDN15, but also provide potential entry points to regulate claudin functions to benefit human health.

## DATA AVAILABILITY STATEMENT

The data are available from the corresponding authors upon request.

## ACKNOWLEDGEMENTS

This work was supported by National Institute of Health R01DK131542 and National Science Foundation grant MCB-1846021.This research is part of the Frontera computing project at the Texas Advanced Computing Center. Frontera is made possible by the National Science Foundation award OAC-1818253.

Le Shen, Fatemeh Khalili-Araghi, and Christopher R. Weber are co-founders and hold stock in Claudyn Biotech.

Author contributions: All authors contributed to writing the manuscript. Fatemeh Khalili-Araghi, Christopher Weber, Le Shen, and Sarah McGuinness, and Shadi Fuladi designed experiments and simulations. Sarah McGuinness, Shadi Fuladi, Sukanya Konar, and Samaneh Sajjadi carried out and analyzed simulations. Pan Li, Ye Li, Mohammed Sidahmed, and Yueying Li carried out and analyzed the *in vitro* experiments.

